# Efficient cardinality estimation for k-mers in large DNA sequencing data sets: k-mer cardinality estimation

**DOI:** 10.1101/056846

**Authors:** Luiz C. Irber, C. Titus Brown

## Abstract

We present an open implementation of the HyperLogLog cardinality estimation sketch for counting fixed-length substrings of DNA strings (“k-mers”).

The HyperLogLog sketch implementation is in C++ with a Python interface, and is distributed as part of the khmer software package. khmer is freely available from https://github.com/dib-lab/khmerunder a BSD License. The features presented here are included in version 1.4 and later.

## Introduction

DNA sequencing technologies have increased in data generation capacity faster than Moore’s Law for more than a decade now, driving the development of new computationalanalysis approaches.

Alon et al. [1996] analyses randomized algorithms for the approximation of “frequency moments” of a sequence using a streaming approach, where items of the sequence are not (or can not be) stored, and are processed sequentially (arrive one by one). KmerStream Melsted and Halldórsson [2014] implemented

A number of probabilistic data structures and algorithms have been developed over the last few years to scale analysis approaches to the rapidly increasing volume of data [Pell et al., 2012],

Here we present an open implementation of the Hyper-LogLog cardinality estimation algorithm, specialized for *k*-length substrings of DNA strings, or *k-mers*. The *HyperLogLog sketch* (HLL) [Flajolet et al., 2008] is a cardinality estimation data structure with constant (and low) memory footprint.

Efficient k-mer cardinality estimation is useful for a variety of purposes, including estimating the memory requirements for de Bruijn graph assemblers [Zerbino and Birney, 2008] and choosing the initial memory allocation for data structures like Bloom filters and Count-Min Sketches ([Zhang et al., 2014]).

## Methods

We implemented HyperLogLog for k-mers in C++ on top of the khmer library. *khmer*[Crusoe et al.] is a library and suite of command line tools for working with DNA sequences. It implements *k*-mer counting, read filtering and graph traversal using probabilistic data structures such as Bloom Filters and Count-Min Sketches. Building on top of khmer leveraged the existing infrastructure (read parsers, package installation, API signatures and some *k*-mer hashing methods).

The *HyperLogLog sketch* (HLL) [Flajolet et al., 2008] estimates the approximate cardinality (*F*_0_ frequency moment) of a set. The HLL is composed of a byte array *M*[1‥*m*] initialized with 0s and a precision value *p*, where

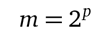

The expected error rate *e* is

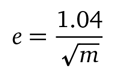

and by modifying *p* we can control the precision of the estimate.

Each position of *M* represents the longest run of zeros found in the *n*th substream, where *n* is an index calculated from the least significant *p* bits of the hashed value:

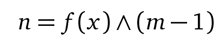

The two basic operations of a sketch are *Add* (or update) and *Merge*. Adding an element *x* involves calculating its hash value using a hash function *f*,

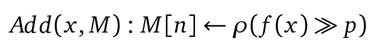

where *ρ*(*h*) is the number of leading zeros in the binary representation of *h*.

The cardinality estimator *E* is the normalized harmonic mean of the estimation on the substream:

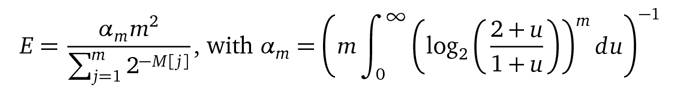

Multiple HLL sketches can be merged by taking the maximum value, element-wise, from every sketch byte-array. For a more detailed description and error bounds analysis, see [Flajolet et al., 2008].

## Implementation details

We chose MurmurHash3 for the hash function because it is one of the fastest non-cryptographic hash functions available and it has a reasonably uniform hash space distribution. Since a *k*-mer is a representation of a substring of a single strand of DNA, the reverse complement on the other strand must also be considered to avoid overcounting. We hash the *k*-mer and its reverse complement individually using MurmurHash3 and create a unique value by doing a binary exclusive-OR on the two hashed values, generating a 128-bit hash value. For compatibility with the current khmer codebase, where hashed values are 64 bits long, we do another binary exclusive-OR over the first and last 64 bits to have a single 64-bit hash value. This procedure is executed len(sequence) − (*k* − 1) times for each sequence in the dataset, where *k* is the desired *k*-mer size.

Our implementation of HLL for multiple processors uses a shared memory model, creating multiple HLL sketches in order to avoid synchronization and locking when adding elements. A master thread processes the input and distributes reads between task threads. After all the reads are consumed the sketches are merged and the final sketch can be used for cardinality estimation of the entire data set. Since sketch sizes are small (16 KiB for a 1% error rate), instantiating additional temporary HLL sketches is a viable tradeoff. One alternative is one HLL shared between threads, with a locking mechanism to isolate the byte array on updates. This would avoid the merge process at the end, but then threads could block on updating the shared structure.

The shared memory model is also a good fit since this is the architecture most potential users have readily available for use. OpenMP is an appropriate choice for the conditions we outlined, and the code compiles to a sequential implementation when OpenMP is not available. We used the OpenMP tasks functionality, since they map well to our problem.

## Benchmarking

All tests were executed on a server hosted by Rackspace. This machine has an Intel Xeon E5-2680 CPU with 20 physical cores (2 sockets) running at 2.80GHz, 60 GB of RAM and a SATA block storage with 200 MB/s transfer rate. During the streaming tests the network transfer rate with the external server was measured to be 10 MB/s.

## Results

### Comparison with exact counters

To test the correctness of the implementation we used two exact cardinality data structures for comparison to our HLL sketch implementation. For exact cardinality we created a Python implementation using the standard library (“collections.Counter”) and another in C++ using the Google sparsehash library. Neither are parallelized. Both implementations are impractical for large cardinalities, so we chose two relatively small datasets from the Sequence Read Archive for benchmarking:

- SRR797943, containing 417 PacBio and 454 sequences with average length 1690 basepairs and totaling 704,951 basepairs, with 670,487 unique *k*-mers. Referred to below as the small dataset.
- SRR1216679, containing 675,236 Illumina reads with average length of 250 bp, 168,809,000 basepairs and 17,510,301 unique *k*-mers. Referred to below as the medium dataset.

Tables 1 and 2 show the runtime, memory consumption, cardinality and the error of the estimate compared to the true cardinality of each dataset. Both exact implementations report the same cardinality and consume similar amounts of memory for each dataset, with the Python implementation taking longer to run. HLL is an order of magnitude faster and consumes a constant amount of memory for both cases, about 6 times less on the small dataset and 200 times less on the medium dataset.

The error in the HLL implementation is close to the ideal upper bound, 1%. The difference can be attributed to MurmurHash3 and our hashing procedure not being perfectly uniform on the hash value space.

**Table 1.** Wall clock time and memory consumption for HLL and two exact cardinality implementations (Python and C++/sparsehash) using the small size dataset.

| Implementation | Time (seconds) | Memory (MB) | Cardinality | Error |
| --- | --- | --- | --- | --- |
| HLL | 0.193 | 13.44 | 670,328 | 0.0002 |
| C++/sparsehash | 3.888 | 77.64 | 670,487 | 0 |
| Python | 4.306 | 83.48 | 670,487 | 0 |

**Table 2.**
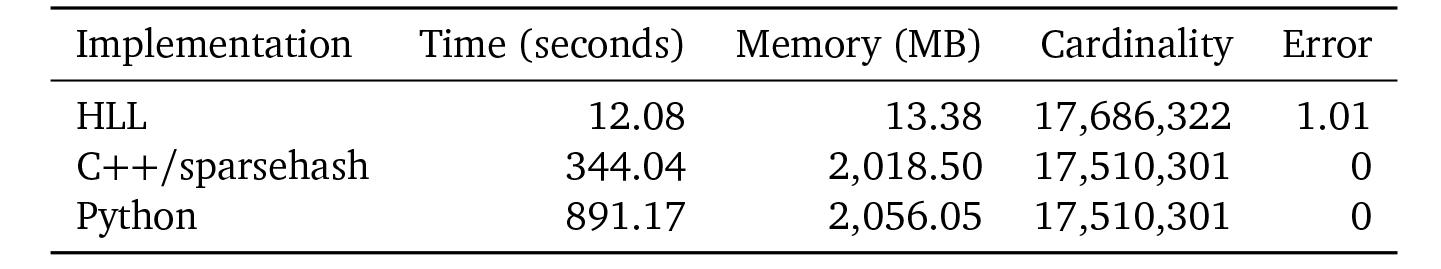
Wall clock time and memory consumption for HLL and two exact cardinality implementations (Python and C++/sparsehash) using the medium size dataset

### Scaling behavior

We chose a larger dataset for examining the scaling performance of our HLL implementation. This larger dataset, SRR1304364, contains 163,379 PacBio sequences with average length 12,934 bp, and 2,113,086,496 basepairs in total.

We examined how our implementation scaled with number of threads. Since hashing is CPU-bound, the problem can be easily parallelized. We ran a simple benchmark to discover the I/O lower bound, using the same input and read parsing infrastructure as the HLL sketch tests, but without performing any kind of processing. Figure 1 shows the results of these tests, where we found 16 threads are needed to saturate I/O on this particular setup, which has 16 physical cores.

**Fig. 1.**
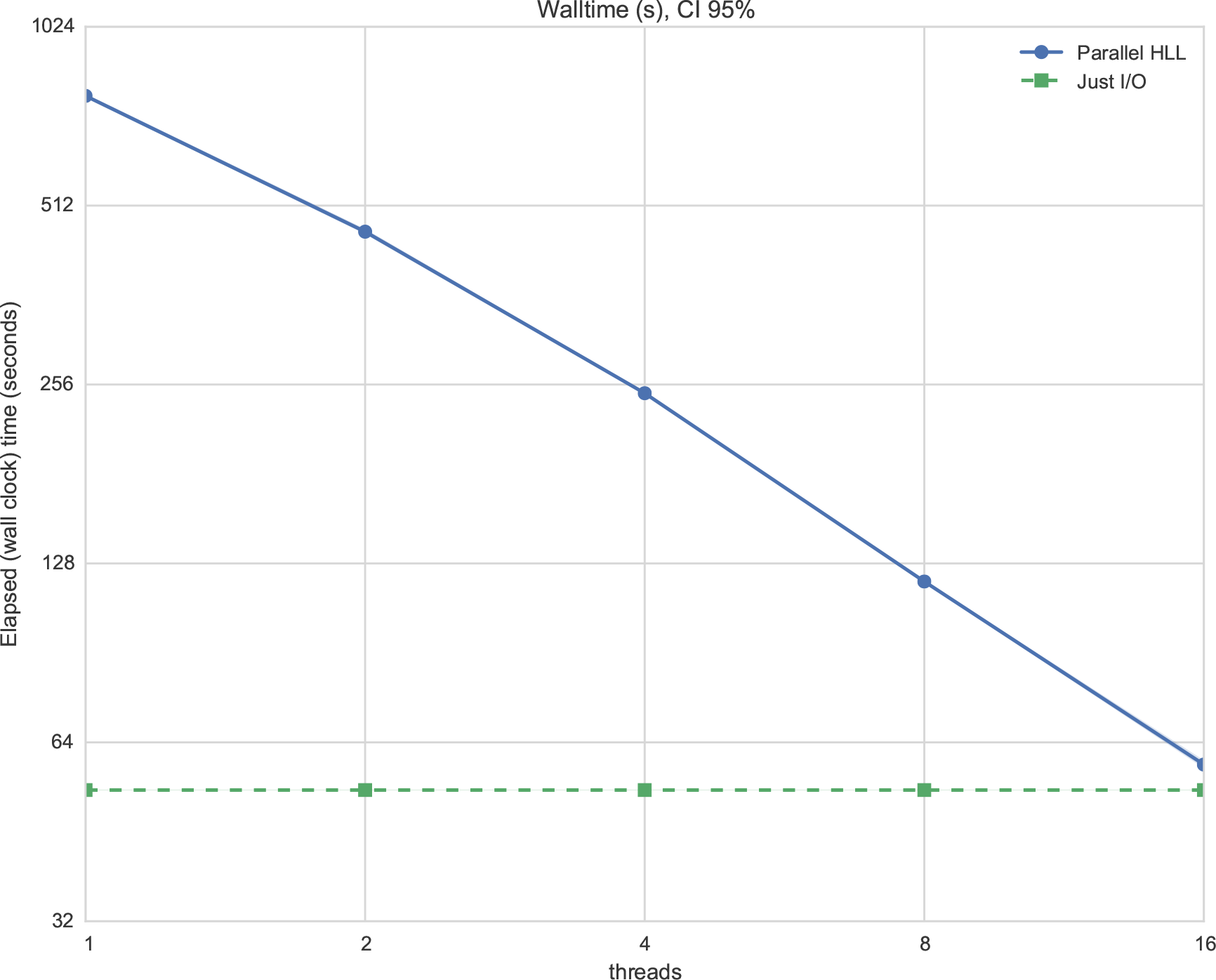
Walltime and lower bound(I/O)

### Streaming

The HyperLogLog sketch is designed for streams of data, and we can take advantage of this property to compose the cardinality estimation capabilities with other pipelines. Here, the overhead of the cardinality counting is minimal with respect to I/O: downloading SRR1304364 and piping it into our HLL implementation adds only 1% to the overall runtime (199.7 ± 9.5 seconds for counting and saving, versus 199.5±13.6 seconds for simply saving the file). Thus our HLL implementation can be used “midstream” to evaluate the effects of streaming lossy compression and error trimming.

## Discussion/Conclusions

We present an open and remixable implementation of the HyperLogLog sketch for cardinality counting of DNA k-mers, written in C++. The implementation scales well to multiple threads, and uses OpenMP for task coordination.

### Author contribution

LCI and CTB conceived the study and designed the experiments. LCI carried out the research, prepared figures and tables and performed the computation work. LCI and CTB analyzed the data and wrote the paper. All authors were involved in the revision of the draft manuscript and have agreed to the final content.

### Grant information

This work was supported by grant 2013–67015–21357 from the United States Department of Agriculture. The funders had no role in study design, data collection and analysis, decision to publish, or preparation of the manuscript.

